# Structural basis for polarized elongation of actin filaments

**DOI:** 10.1101/2020.06.02.129940

**Authors:** Vilmos Zsolnay, Harshwardhan H. Katkar, Steven Z. Chou, Thomas D. Pollard, Gregory A. Voth

## Abstract

Actin filaments elongate and shorten much faster at their barbed end than their pointed end, but the molecular basis of this difference has not been understood. We use all-atom molecular dynamics simulations to investigate the properties of subunits at both ends of the filament. The terminal subunits tend towards conformations that resemble actin monomers in solution, while contacts with neighboring subunits progressively flatten the conformation of internal subunits. At the barbed end the terminal subunit is loosely tethered by its DNase-1 loop to the third subunit, because its monomer-like conformation precludes stabilizing contacts with the penultimate subunit. The motions of the terminal subunit make the partially flattened penultimate subunit accessible for binding monomers. At the pointed end, unique contacts between the penultimate and terminal subunits are consistent with existing cryo-EM maps, limit binding to incoming monomers, and flatten the terminal subunit, which likely promotes ATP hydrolysis and rapid phosphate release. These structures explain the distinct polymerization kinetics of the two ends.

**Significance Statement:** Eukaryotic cells utilize actin filaments to move, change shape, divide, and transport cargo. Decades of experiments have established that actin filaments elongate and shorten significantly faster from one end than the other, but the underlying mechanism for this asymmetry has not been explained. We used molecular dynamics simulations to investigate the structures of the actin filament ends in the ATP, ADP plus *γ*-phosphate, and ADP nucleotide states. We characterize the structures of actin subunits at both ends of the filament, explain the mechanisms leading to these differences, and connect the divergent structural properties of the two ends to their distinct polymerization rate constants.

## Introduction

Actin is one of the most abundant proteins in eukaryotic cells. By polymerizing into filaments, actin and its associated proteins form polymer networks that drive cell motility, division and morphological change. Critical to these functions, the elongation rate at one end, the ‘barbed end’, is much faster than the other ‘pointed end’. Although this asymmetric growth has been known for decades (1–3), the underlying basis that gives rise to these vastly different kinetic rates has not been understood. The lack of structural data of subunits at the filament ends has limited this understanding.

Monomeric actin may be crystallized, which has allowed the determination of many high-resolution crystal structures of the actin monomer (4–7). Actin filaments have not been crystallized, and so X-ray fiber diffraction (8) and cryo-electron microscopy (9, 10) have been employed to determine actin filament structures at resolutions up to 3.1 Å. These reconstructions have revealed that the primary conformational change between monomeric and filamentous actin is a flattening of the actin molecule. This flattening can be quantified by measuring the dihedral angle made by actin’s four subdomains (Fig. 1A), which is around −18° for actin monomers and around −3° for subunits within filaments (Fig. 1B). The flattened conformation increases the hydrolysis rate of actin’s bound ATP by a factor of 10^4^ (11–14).

**Fig. 1.**
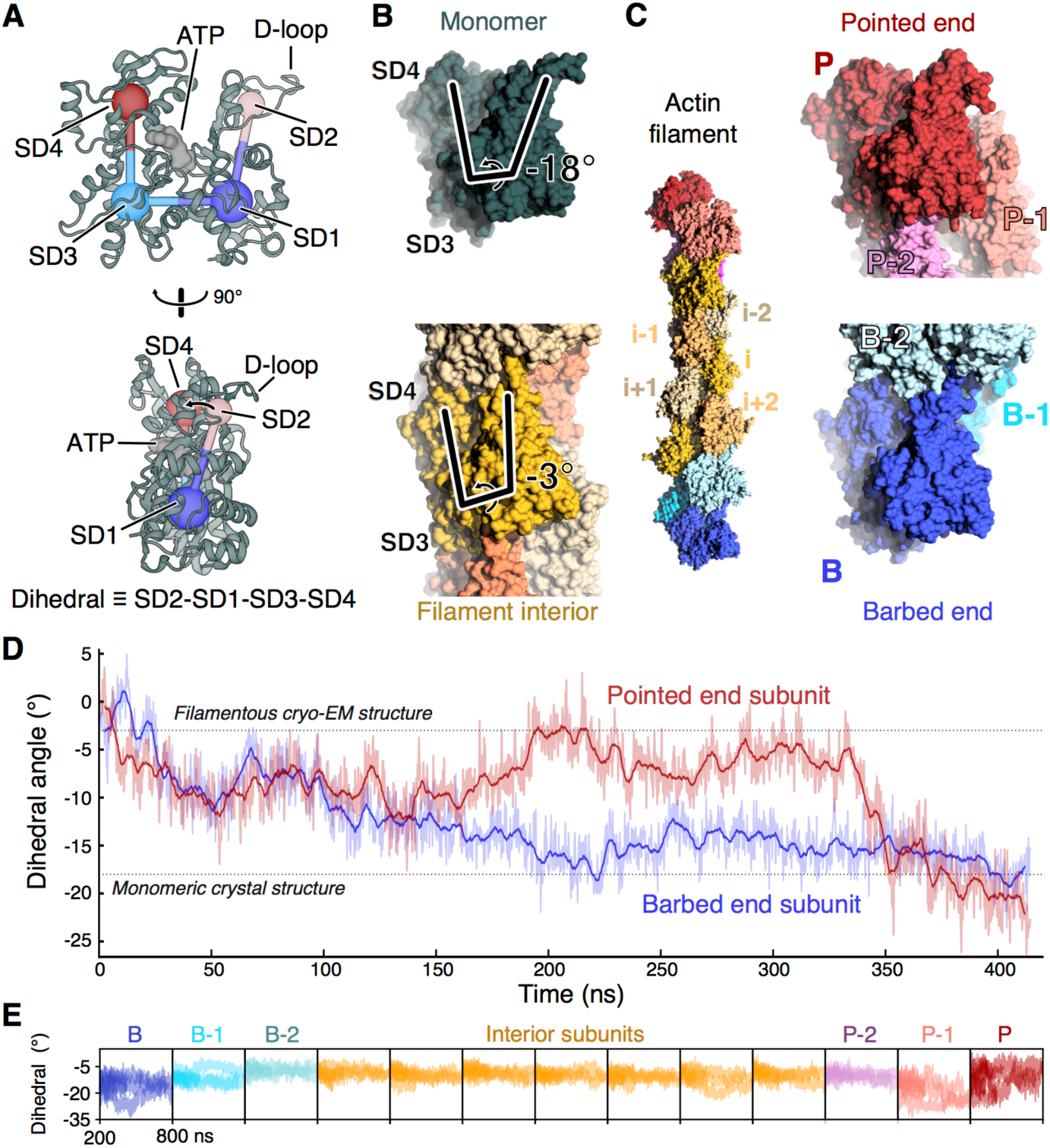
The terminal subunits of actin filaments take on monomer-like conformations. (**A**) Two orientations of a ribbon diagram of the ATP-bound actin monomer with labels on the four subdomains, SD1-4. The dihedral angle defined by SD2-SD1-SD3-SD4 characterizes the primary structural transition between monomeric actin and polymerized, filamentous actin. The flexible D-loop in SD2 forms stabilizing contacts between subunits. Protein data bank (PDB) entry 3TU5. (**B**) Side views of space-filling models of monomeric actin (dark green, PDB 3TU5) and a subunit within a filament (yellow, PDB 6JDM). The lines highlight the difference in the dihedral angle between actin monomers (−18°) and a subunit in a filament (−3°). The models are aligned by subdomains 3 and 4. (**C**) Left: Space-filling model of the final frame of a 415-ns molecular dynamics simulation of an ATP-actin 13-mer. Note that the terminal subunits at the barbed end (B, blue) and pointed end (P, red) lack two neighboring subunits that form contacts between interior subunits. The penultimate subunits B-1 and P-1 each lack one neighboring subunit. Right: Zoom in on the filament ends showing the terminal subunits B and P in conformations closely resembling monomers. Subunits B and P are aligned relative to each other by subdomains 3 and 4. (**D**) Time course of changes of the dihedral angle of the terminal P and B subunits during a 415-ns simulation of the ATP-actin filament, in which all subunits started in the conformation of interior subunits (PDB 6DJM). During the simulation, both terminal subunits transitioned spontaneously from the filamentous conformation toward the monomer conformation. Raw data (light) and 4-ns moving average (dark) are shown. Data and snapshots are from simulation 3 (Table S1). (E) Time courses of the dihedral angles of each subunit in the 13-mer actin filaments for times greater than 200 ns. Seven runs are shown (simulations 1-7 (Table S1)). Subunits at the ends sample a wider range of conformations than the flattened internal subunits (yellow).

Acquiring structures of subunits at the filament ends is a much greater challenge. So far, only the structure of the pointed end has been solved at a resolution of 22.9 Å (15). In the absence of structural data, it has largely been assumed that subunits at the filament ends adopt the flattened conformation of interior subunits.

Here we use all-atom molecular dynamics (MD) simulations of an actin filament to investigate the equilibrium conformations of subunits at the filament ends. Specifically, we constructed actin 13-mers from structures of interior subunits for each of the ATP, ADP-P_i_, and ADP nucleotide states (Fig. 1C, left). Our simulations show that the barbed end and pointed end subunits adopt distinct equilibrium conformations, which lead to meaningful differences in the contacts between neighboring subunits. These distinct structural properties lead to natural explanations for the observed differences in actin elongation kinetics of the two filament ends.

## Results

### Subunits at filament ends transition to monomer-like conformations

In all-atom MD simulations of actin filaments, subunits at both ends spontaneously transitioned from the flattened conformation found in the middle of filaments to conformations with larger negative dihedral angles that resemble free actin monomers (Fig. 1, C and D). The relaxation to a large dihedral angle was generally less pronounced for subunits further from the ends (Fig. 1E), suggesting that the transition to the flattened conformation does not occur discretely when a subunit incorporates into the filament, but rather, occurs as contacts formed by addition of new monomers facilitate a gradual transition from the monomeric conformation to the structure of interior subunits.

### ATP Barbed End

A major consequence of monomer-like conformations of barbed end subunits is that a monomeric subunit cannot make the full set of contacts that connect subunits i and i-1 in the interior of the filament. This is due to the fact that subdomain 2 and subdomain 4 of subunit i must be roughly planar (i.e., subunit i must be flat) to make contacts with subunits i-1 and i-2 simultaneously. At the ATP-bound barbed end, subunit B takes on a monomeric conformation (Fig. 2A, top), and therefore cannot make one or more contacts with subunits B-1 or B-2.

**Fig. 2.**
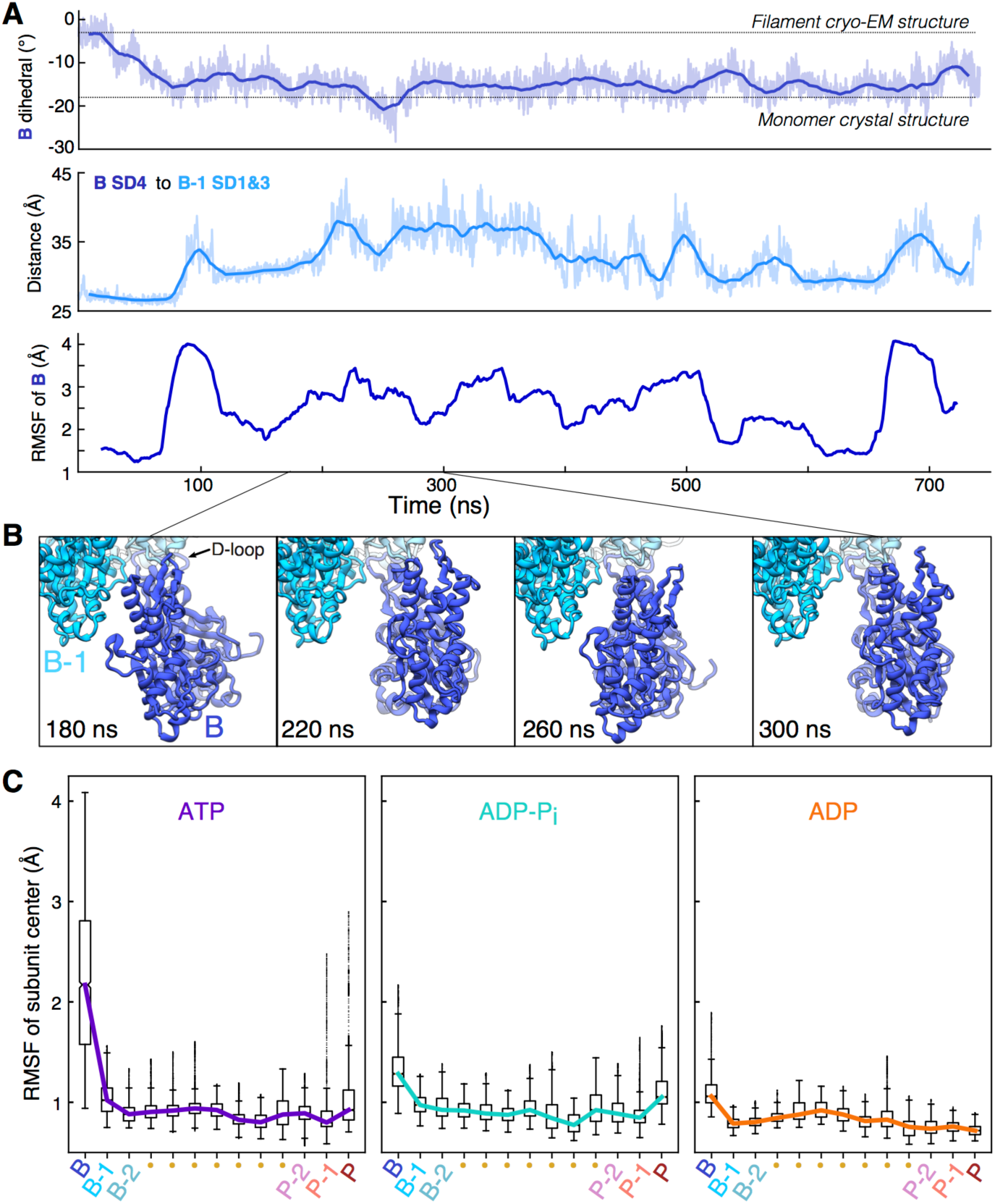
The transition of the ATP barbed end subunit to a monomeric dihedral angle is associated with loss of lateral subunit contacts, while remaining tethered to the barbed end by its D-loop. (**A**) Time course of changes during a 742-ns MD simulation of an ATP-actin filament. The top and middle figures show raw data (light) and 20-ns moving averages (dark). Top: The dihedral angle of barbed end subunit B spontaneously increases as the subunit transitions to a monomeric conformation. Middle: The distance between centers of mass of subdomain 4 of subunit B and subdomains 1 and 2 of subunit B-1 increase as subunit B transitions to a monomer-like structure. Bottom: The root-mean-square fluctuation (RMSF in 40-ns windows) center of mass of subunit B increased in step with the transition to a monomer-like conformation and the loss of inter-subunit contacts. (**B**) Ribbon diagrams of the terminal subunit B at 40-ns intervals showing movements around the tether formed by the D-loop of subunit B stably bound to subunit B-2. Data are from simulation 1. (**C**) Boxplots of the center of mass RMSF for each subunit in the 13-mer reveal gradients of fluctuations from both ends the filament. Barbed end subunits with bound ATP fluctuate the most and those with ADP fluctuate the least. Data comes from times greater than 200 ns in simulations 1-3 for ATP, simulations 4-6 for ADP-P_i_, and simulation 7 for ADP (Table S1).

In keeping with the transition of subunit B to a large dihedral angle, subunit B lost its contacts with subunit B-1, and the distance between subdomain 4 of subunit B and subdomains 1 and 3 of subunit B-1 increased from ∼26 to ∼37 Å (Fig. 2A). The loss of contacts between them (Fig. S1) led to pronounced dynamics of subunit B, which only maintained stabilizing attachments to subunit B-2 through its D-loop in subdomain 2 (Fig. 2B, Movie S1).

Calculations of the root-mean-squared fluctuation of subunit B’s center of mass relative to neighboring subunits B-1 and B-2 (see Methods) revealed a large and sustained increase in the motion of subunit B following the loss of lateral contact with B-1 (Fig. 2A, bottom). Despite these fluctuations, the D-loop of B remained securely associated with B-2 throughout the entire simulation, and complete dissociation of subunit B from the filament appeared unlikely on the simulation timescale. Fluctuations of the barbed end subunit were consistently higher than elsewhere in the filament.

The nucleotide bound to the barbed end subunit influenced its dynamic fluctuations (Fig. 2C). The ATP-bound barbed end subunit fluctuated most, with a RMSF interquartile range (IQR) spanning 1.6 to 2.8 Å. In contrast, the ADP-bound barbed end subunit was least dynamic with an IQR spanning 1.0 to 1.2 Å. Subunit B in this nucleotide state fluctuated less because it relaxed to a lesser dihedral angle (∼-12°), allowing it to maintain its lateral contacts with subunit B-1. The RMSF of subunit B in the ADP-P_i_ state had an intermediate IQR spanning 1.2 Å to 1.5 Å, which resulted from generally weakened lateral contacts. However, these associations were not so weak that full separation between subunits B and B-1 occurred consistently, as in simulations of ATP-bound filaments.

### ADP Pointed End

The tendency for terminal subunits to take on monomer-like conformations led to changes at the pointed end (Fig. 3) almost opposite to those at the barbed end. In simulations of ADP-actin filaments, the penultimate pointed end subunit, P-1, which lacks an i-2 neighbor to secure its subdomain 2, initially sampled a broad range of dihedral angles. Wide dihedral angles brought residues in subdomain 2 of subunit P-1 close to subunit P, the terminal subunit at the pointed end (Movie S2). These contacts allowed attractive interactions to form between the neighboring subunits P and P-1. Of the interacting residues, the most stable contacts were between the sidechains of R62 of subunit P-1 with E270 of subunit P (Fig. 3A, left). In addition, the sidechain of R39 of subunit P-1 formed a hydrogen-bond network with oxygens of several nearby residues (Fig. S2). As a result, subdomain 2 of P-1, including the structured helix composed of residues 55-64, shifted towards subunit P and was secured in a monomer-like conformation (Fig. 3A and Movie S2). Furthermore, the D-loop of P-1 formed a long-lasting association with subunit P (Fig. 3A, right and Movie S2), although the specific residue-residue contacts varied over time.

**Fig. 3.**
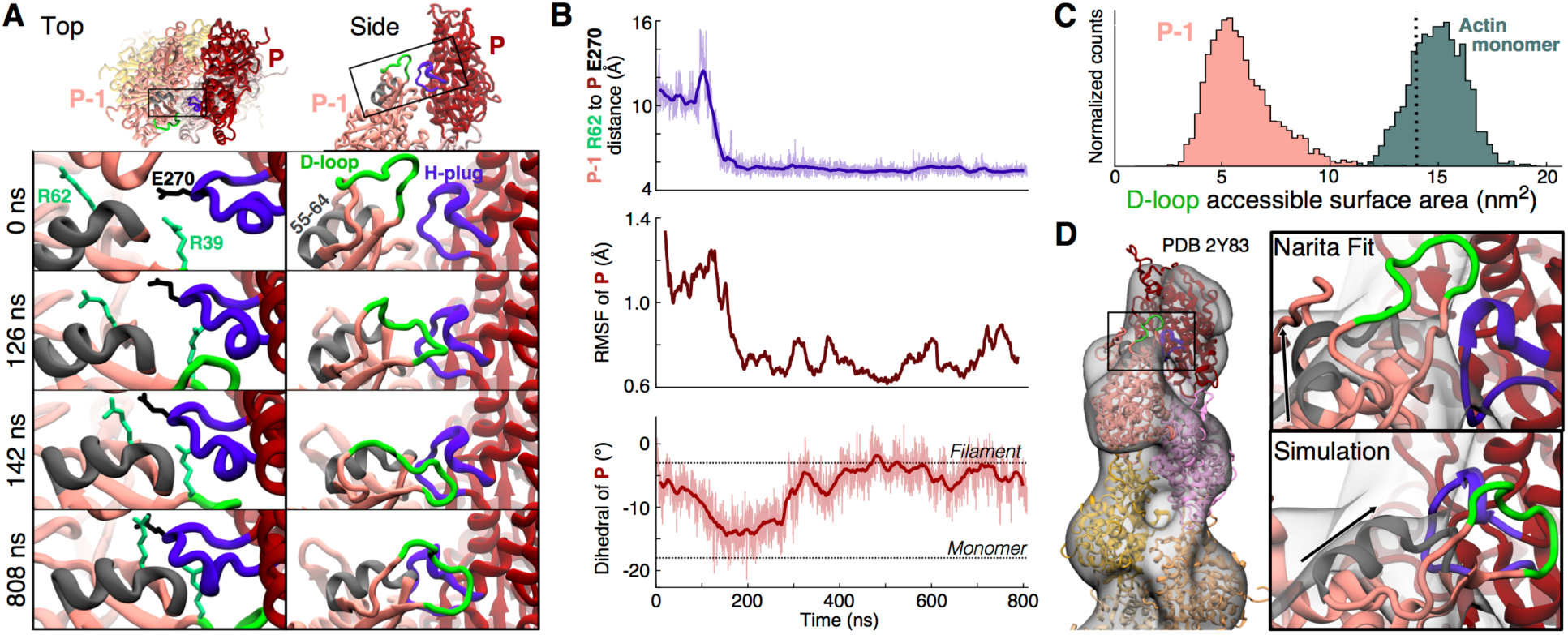
Transition of the ADP pointed end subunit P-1 to a monomeric dihedral angle gives rise to stabilizing contacts with subunit P that flatten P and sequester the D-loop of P-1. (**A**) Ribbon diagrams from two views of the pointed end of an ADP-actin filament showing unique contacts formed during MD simulations between subdomain 2 of subunit P-1 and subdomains 3-4 of subunit P. The four panels in each column show selected time points. Left: looking down the filament axis at the pointed end, R39 and R62 of subunit P-1 (foam green) contact the hydrophobic plug (violet) of subunit P (E270 shown in black). These long-lasting interactions stabilize a shift in the position of a helix composed of residues 55-64 (gray) of subunit P-1. Right: a side view depicts the progressive shift of subdomain 2 of P-1 toward subunit P resulting in the D-loop of subunit P-1 (green) forming a stable, lateral contact with subunit P. (**B**) Time course of the formation of the connection between subunits P and P-1 that stabilizes and flattens the pointed end. Raw data (light) and 20-ns moving averages (dark) are shown. Top: Distance between the C_α_ atoms of R62 of subunit P-1 and E270 in the H-plug of subunit P. Middle: The 40-ns RMSF of subunit P’s center of mass drops in step with the formation of contacts with P-1. Bottom: The dihedral angle of subunit P transitions toward the monomer conformation, then back to flattened structure after forming contacts with P-1 (**C**) Histograms of the accessible surface areas at a radius of 7 Å around the D-loops of subunit P-1 and actin monomers. The dotted line marks the value assuming P-1 takes on the flattened conformation of interior subunits (PDB 6DJO). Data and snapshots are from simulation 7 for P-1 and simulation 9 for G-actin (Table S1). (**D**) Comparison of pointed end models with the 3D reconstruction of the ADP-actin pointed end by Narita et al (2011). Left: Narita model with P-1 in yellow and P in orange. Right: Enlargement of the bridge-like density connecting P-1 and P in the model. Our simulated ADP structure (bottom) places the structured alpha-helix of subunit P-1 (black arrow) and H-plug of P within the bridge density with the D-loop stably attached to subunit P.

The fluctuations of subunit P decreased significantly after this new association with subunit P-1 (Fig. 3B) and remained lower than the average fluctuations of interior subunits (∼0.9 Å) (Fig. 2C) in spite of lacking two neighboring subunits, i-1 and i-2. Consequently, the pointed end subunits were very rigid with respect to each other in the ADP-state, in contrast to the loosely-attached and flexible ATP-bound barbed end. In addition, the formation of contact with P-1 reversed the transition of subunit P to the monomeric conformation (Fig. 3B, bottom), supporting the concept that monomeric conformations of terminal subunits arise from a lack of contact with neighboring subunits.

A further consequence of the interaction between subunits P-1 and P is that the D-loop of P-1 is far less available to form other protein-protein interactions—namely, bind the barbed end of a nearby free actin monomer—due to its close association with subunit P. To quantify this effect, we calculated the accessible surface area (ASA) around the D-loop at a probe radius of 7 Å (Fig. 3C). Throughout the simulation, the D-loop of P-1 was greatly occluded, with the most common ASA being ∼5.25 nm^2^. For comparison, the ASA assuming subunit P-1 were in the conformation of an interior subunit is ∼14 nm^2^ (Fig. 3C dotted line, PDB 6DJO). More consequential for polymerization rate kinetics, however, is the comparison with the D-loop of a free actin monomer. At the barbed end of the filament, it is the monomer’s D-loop that participates in the connection with subunit B-1. Strikingly, the accessibility of the monomer’s D-loop is nearly three times that of subunit P-1, with a most likely ASA of ∼15.25 nm^2^. Furthermore, the two distributions hardly overlap (<1% intersection). This disparity implies that the stabilizing D-loop contact between a free monomer and a filament end forms much more readily at the barbed end than at the pointed end. Additionally, it is likely that the D-loop of subunit P-1 must lose its association with subunit P in order for an actin monomer to become fully incorporated at the pointed end.

The connections between subunits P and P-1 formed readily (within 200 ns) in both ADP simulations, including one which was initialized with the D-loops in helical conformations, but did not form in our original ∼350-ns ATP and ADP-P_i_ simulations. However, in one extended simulation of the ADP-P_i_ filament, the connection observed between subunits P and P-1 in ADP-filaments emerged after ∼580 ns and remained thereafter. The extended ATP replica reached 742 ns and the connection did not form. However, subunit P-1 assumed a large dihedral angle, which was a precursor for this connection to form in the other nucleotide states. Therefore, we propose that the contacts connecting subunit P-1 to subunit P described above are likely to occur in the ADP and ADP-P_i_ states, but are less probable in the ATP state, though likely not forbidden. It should be noted that because phosphate dissociates rapidly from pointed end subunits (16), ATP and ADP-P_i_ will occupy pointed end subunits only transiently, favoring the interactions between subunits P and P-1.

The stable connection that forms at the pointed end between subdomain 2 of subunit P-1 and subdomains 3 and 4 of subunit P in our simulations bears striking resemblance to electron potential maps of the pointed end of ADP-bound filaments (15). Although the resolution was limited to 22.9 Å, that reconstruction revealed a bridge-like density connecting subunit P-1 to subunit P. The authors proposed a model whereby the D-loop of subunit P-1 and the hydrophobic plug of subunit P partially occupy the bridge density by forming lateral contact. However, in their proposed fit, P-1 is mostly in the conformation of a flattened subunit, and so the alpha-helix of subdomain 2 does not match the bridge density well (Fig. 3D, right). In contrast, the connection between subunits P and P-1 that emerged in our MD simulations forms after a significant shift of subdomain 2 of P-1 towards subunit P. Because of this shift, the conformation emerging from our simulations fits better into the bridge density (Fig. 3D, right). Additionally, terminal subunit P in the electron potential map appears to be flattened, in agreement with our simulations of the ADP-pointed end.

## Discussion

We find the mechanism that underlies the distinct structures at the two ends of actin filaments is in fact the same: subunits with limited contacts with neighboring subunits are free to adopt the conformation of monomers with large dihedral angles. Monomeric actin without neighboring subunits has a dihedral angle of ∼-18°, whereas interior subunits within filaments with four neighboring subunits assume a flat dihedral angle around −3°. Therefore, terminal subunits, which lack one or two neighbors, may take on intermediate dihedral angles (Figs. 1E and 4). The tendency for terminal subunits to assume large dihedral angles leads to unique structures at the two filament ends, which explain their different elongation kinetics, as detailed below.

**Fig. 4.**
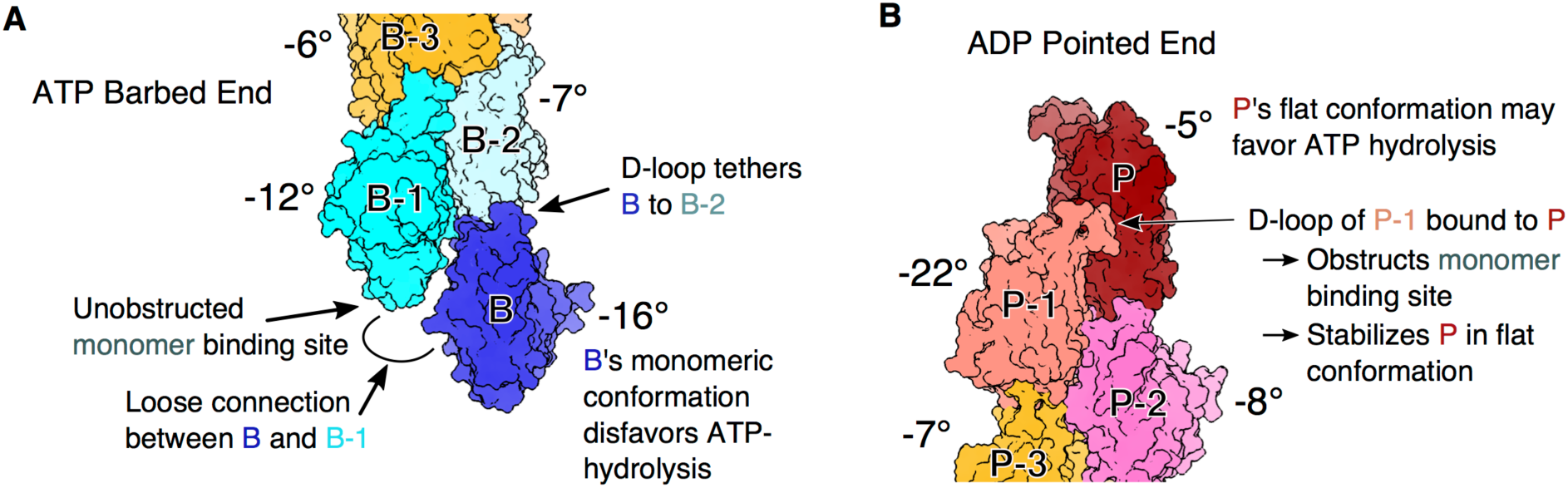
Structural basis for polarized actin filament elongation kinetics. Space-filling models of subunits at the ATP barbed end and ADP pointed end. Median dihedral angles are shown next to each subunit. (**A**) Subunits incorporated at the barbed end are progressively flattened as they make contacts with newly polymerized subunits, so ATP hydrolysis is more likely in interior subunits. Subunit B’s monomeric conformation hinders stable contact with subunit B-1, leaving the D-loop of subunit B tethered to subunit B-2 and the monomer-binding site on subunit B-1 unobstructed. (**B**) Unique contacts formed between subdomain 2 of subunit P-1 and subunit P greatly limit interactions of the D-loop of P-1 with monomers and stabilize subunit P in a flattened conformation that may favor hydrolysis of the bound ATP.

Our simulations show that flattening generally progresses as subunits acquire neighbors at the barbed end (Fig. 4A). This progression to a flat conformation appears to be gradual and cooperative. Each additional lateral contact further stabilizes the flat conformation and strengthens connections along the short-pitch helix, as predicted from reconstructions of interior subunits (10). Pointed end subunits P and P-1 are different. The monomeric conformation of subunit P-1 facilitates the formation of a unique lateral contact with subunit P, which is linked to flattening of this terminal subunit in spite of having only two neighbors (Fig. 3B and 4B). The lateral contact between subunits P and P-1 is not observed elsewhere in filaments but stabilizes the unique conformation of pointed ends originally observed by Narita et al. (15).

### Nucleotide states of terminal subunits

Flattening actin subunits increases ATP-hydrolysis 10^4^-fold (11–14), so the progressive flattening of terminal subunits must influence the nucleotide state of subunits at both filament ends. Our simulations suggest that ATP-hydrolysis is delayed at the barbed end until subunits have been flattened by contacts with four neighbors (i.e., position B-2 and deeper in the filament). Thus, the terminal barbed end subunits likely have bound ATP.

At the pointed end, the unique lateral contact made by penultimate subunit P-1 flattens terminal subunit P (Fig. 3B, bottom), which should favor ATP hydrolysis. If subunit addition is slow, this temporary flattening may result in the hydrolysis of ATP bound to most new subunits added to the pointed end. For example, at steady state with about 0.1 µM free ATP-actin monomers, the association rate is ∼0.1 subunit per second and the hydrolysis rate within the filament is 0.3 s^-1^. In simulations of ADP and ADP-Pi filaments, the R177 gate on the phosphate release channel (10) is mostly open on subunit P and opens for subunit P-1 in step with the formation of the contact between subunits P and P-1. Open gates on these two subunits may explain why phosphate dissociates rapidly at the pointed end compared with the barbed end and interior subunits (16). Rapid dissociation of phosphate from subunit P-1 would favor formation of the contact with subunit P.

These features suggest a cooperative mechanism for ATP hydrolysis and phosphate dissociation by pointed end subunits. Specifically, assuming newly incorporated subunit P arrives with bound ATP, having ADP bound to subunit P-1 makes more likely the formation of the lateral connection and flattening of subunit P, favoring the hydrolysis of its bound ATP. On the other hand, with high concentrations of ATP-actin monomers, subunit addition would exceed the rate of ATP hydrolysis on subunit P, so subunit P-1 would also have bound ATP, reducing the chance that it would form the connection with subunit P, leaving P in its monomeric conformation and less likely to hydrolyze its bound ATP until it is deeper in the filament.

### Elongation at barbed ends

Elongation at barbed ends is a diffusion-limited reaction with a high probability that random collisions with incoming subunits result in binding (17). The dependence of the elongation rate on the solution viscosity demonstrated that the reaction is diffusion limited, and the calculated collision rate constant is only 50 times higher than the observed rate constant, giving a very high orientation factor (probability of binding) of 0.02 in the Smoluchowski equation. The high degree of freedom of barbed end subunits revealed by our simulations may contribute to making these reactions highly favorable in three ways.

First, monomer binding to subunit B-1 is likely enhanced by the separation of subunit B from subunit B-1 (Movie S1), which opens up the initial binding site for the D-loop on subunit B-1 (Fig. 4A). This mobility of subunit B results from its monomeric conformation, which precludes stable lateral contacts with B-1. As a result, subunit B is only attached to the filament through association of its D-loop with subunit B-2. Despite this single attachment point, the large buried surface area of the D-loop connection (10) tethers terminal subunit B to the filament end throughout energetic fluctuations. Thus, the D-loop association alone suffices to bind an incoming subunit B+1 to the barbed end.

Second, partial flattening of subunit B-1 may favor binding of the D-loop of an incoming actin monomer. Flattening of interior subunits opens a hydrophobic pocket for binding M44 from the D-loop of the next subunit along the long-pitch helix (10). In our simulations, this pocket on subunit B-1 was generally closed even though the subunit relaxed to an intermediate dihedral angle around −11°. However, subunit B-1 spent ∼12% of the time with dihedral angles flatter than −6°, which may favor transient opening of the pocket for M44.

Third, the simulations also show that tethered subunit B spontaneously forms lateral associations with subunit B-1. For example, the extended ATP-filament simulation revealed that after losing the starting contacts between subunits B and B-1, subunit B transiently visited a state that reestablished some of the contacts with subunit B-1 ∼500 ns later. These contacts lasted ∼50 ns before dissociating again (Movie S1, Fig. S1). It is reasonable to believe that this state would be visited periodically in a longer simulation. If this were true, then every half microsecond or so, the tethered subunit B will spontaneously approach a state where lateral contacts with subunit B-1 form easily. Therefore, when incoming subunit B+1 makes contact with B-1, subunit B is poised to quickly establish lateral contact with B-1 and participate in the next polymerization reaction (i.e., accept contact from subunit B+2).

### Elongation at pointed ends

At the pointed end, elongation is slow and not diffusion-limited (17). Our simulations show that the monomeric conformation of subunit P-1 allows its D-loop to attach to subunit P (Fig. 3A), which reduces the probability that the D-loop can bind an incoming monomer. It is likely that this association is essential for an incoming subunit to be incorporated as P+1, because the D-loop connection appears to be the strongest stabilizing contact between subunits at the barbed end (Fig. 2B) and in the filament interior (10). Therefore, unbinding of the D-loop of subunit P-1 from subunit P is a pointed-end-specific step that limits the rate of elongation, as predicted by Narita et al. (15). Compromising the reaction further, the pointed end connection formed most readily in the ADP-nucleotide state, which is also the most probable nucleotide state for subunits P and P-1 given that phosphate dissociates rapidly from pointed end subunits (16) and P is flattened by subunit P-1 (Fig. 3B). Additionally, the barbed end of the free monomer does not have a favorable conformation to bind the D-loop of subunit P-1, which could also limit pointed end elongation.

### Methodological considerations

All atomistic MD studies are limited to finite timescales, generally in the hundreds of nanoseconds to microsecond range, so we cannot rule out that other properties would emerge in longer simulations. In particular, we began the simulations with all subunits in the conformations of interior subunits. If this initial structure were to be very far from the true equilibrium conformations of the ends, then reaching the preferred orientations may not presently be possible in all-atom MD simulation. However, the close match to existing structural data of the pointed end (15) suggests that our all-atom approach adequately sampled conformational space in order to reach equilibrium distributions. Furthermore, recent coarse-grained Monte Carlo simulations of an actin monomer associating with an actin dimer reported high-probability barbed end conformations similar to ones sampled in our all-atom simulations (18).

### New mechanisms for actin binding proteins

In addition to establishing a structural basis for the great differences in elongation rate constants at the two ends of bare actin filaments, our simulations provide a new framework for understanding the behavior of actin binding proteins that interact with terminal subunits. As an example, profilin has a high affinity for the barbed end of actin monomers, and a weak affinity for subunits in the flattened conformation. Concentrations of profilin that saturate barbed ends slow elongation (19, 20). Previously, it was assumed that the terminal subunits at the barbed end are flattened, but our simulations demonstrate that terminal subunit B has a monomeric conformation and subunit B-1 is only partially flattened so both may be able to bind profilin. Additionally, the freedom of subunit B reduces steric clashes when both subunits B and B-1 are bound to profilin. These insights lead to a straightforward mechanism by which the FH2 domains of formins can promote the dissociation of profilin from subunit B: contacts with the FH2-domain may flatten subunit B, lowering its affinity for profilin.

## Materials and Methods

### System setup for molecular dynamics simulations

We constructed actin filaments in the ATP, ADP-P_i_, and ADP-bound nucleotide states by patterning 13 of the corresponding F-actin structures reported in (10) (PDB: 6DJM, 6DJN, 6DJO) and using the accompanying rise and twist values. For PDB 6DJM, we substituted AMPPNP for ATP. We included waters in the catalytic center from previously-equilibrated actin filament simulations, and solvated the filament structure using VMD (21) plugin auto-solvate such that at least 11 Å of TIP3P water solvated each direction. We used the auto-ionize plugin to reach a charge-neutral system with a KCl concentration of 100 mM. Periodic boundaries were active across each box dimension, meaning at least 22 Å of buffer separated the protein from its periodic image. The CHARMM27 force field with CMAP correction was used (22).

Each of the systems were sequentially energy minimized in NAMD version 2.11 (23) for 200 ps using a 2-fs timestep under each of the following harmonic constraint selections (k = 10 kcal•mol^-1^•Å^-2^): (a) everything except buffer; (b) everything except buffer and protein sidechains; (c) only the bound nucleotide and Mg^2+^; (d) only Mg^2+^. This was followed by a heating protocol lasting 1 ns from 0 to 310K under constraint selection 1. The systems were then equilibrated with constraint selection (a) being progressively weakened by a factor of 2 every 400 ps for 5 iterations (i.e., k = 10, 5, 2.5, 1.25, 0.625 kcal•mol^-1^•Å^-2^) and followed by a final constrained equilibration with k = 0.1 kcal•mol^-1^•Å^-2^ for 1 ns. The systems were equilibrated without constraints for 6 ns. Time courses include data immediately following equilibration, which illustrates transitions. However, aggregate data analysis included data following an additional 200 ns of simulation, which represented the period after major structural transitions occurred.

Production runs were performed on NSF XSEDE supercomputers using the final frame of the equilibration protocol as an initial structure. These were performed using GROMACS version 2018.3 (24) with the leap-frog integrator in the isothermal-isobaric (constant *NPT*) ensemble using Parinello-Rahman pressure coupling and v-rescale temperature coupling. Electrostatic interactions were calculated using the particle mesh Ewald sum method with a cutoff of 1.2 nm.

Set up of the simulation of monomeric actin (PDB 1NWK) (4) was performed using the same protocol. Production runs were performed in NAMD on group-owned compute nodes (see Table S1 for initial structures and MD run times).

Construction of simulation 8 (ADP with folded D-loop; Table S1) used an early structure of the ADP subunit (25), in which residues 36-58 were replaced by corresponding residues in simulations from reference (26) that had the D-loop in a helical conformation. The rest of the construction, minimization, and production protocol was the same as with the other simulations.

### Analysis protocols for molecular dynamics simulations

For the calculation of the dihedral angle, actin was divided into four subdomains following standard residue assignments: Subdomain 1 (SD1): residue 1 to 32, 70 to 144, 338 to 375; SD2: residue 33 to 69; SD3: residue 145 to 180, 270 to 337; SD4: residue 181 to 269. The center of mass of the C_α_ atoms of each subdomain was calculated and these positions were used to compute a dihedral angle SD2-SD1-SD3-SD4.

We reported a measurement of subunit-level root-mean-squared fluctuation (RMSF) relative to neighboring subunits. To calculate this metric for subunit i, we aligned the simulation to the C_α_ atoms of two of the subunit’s neighbors, either (i+1, i+2) or (i-1, i-2) depending on positioning in the filament. We then mapped subunit i into a single coarse-grained (CG) bead using the C_α_ center of mass of subunit i for its position. Finally, we calculated the root-mean-square fluctuation (RMSF) of the CG bead throughout 40-ns windows for the simulation.

To quantify the occlusion of the D-loop, we calculated the accessible surface area (ASA). This analysis draws a surface one probe radius around the D-loop and calculates how much of it is blocked by something other than water or ions. ASA calculations are commonly performed in the context of solvent exposure, where the probe radius is chosen to approximate the radius of a water molecule (1.4 Å). In our study, the relevant context is the D-loop’s availability to form new protein-protein interactions, so the probe radius was chosen to approximate the distance at which sidechains interact with each other. To account for the many ways in which sidechains are able to interact, we chose a 7 Å probe radius.

## Acknowledgments

This work was supported through the Department of Defense Army Research Office through a Multidisciplinary Research Initiative Grant W911NF1410403 (V.Z., H.H.K., and G.A.V.) and by the National Institute of General Medical Sciences of the National Institutes of Health under award number R01GM026338 (S.Z.C. and T.D.P.). The content is solely the responsibility of the authors and does not necessarily represent the official views of the National Institutes of Health. Simulations were performed on National Science Foundation XSEDE resources at the Texas Advanced Computing Center (TACC). The authors thank Prof. David R. Kovar and Dr. Sriramvignesh Mani for helpful discussions.

## Notes

### Competing Interest Statement

The authors have declared no competing interest.

