## Supporting Information for "Structural basis for polarized elongation of actin filaments"

**This PDF file includes:**

Legends for Movies S1 and S2
Figures S1 and S2
Table S1

**Other supplementary materials for this manuscript include the following:**

Movies S1 and S2

**Movie S1. The ATP-bound barbed end subunit B breaks lateral contact and fluctuates dynamically.** A movie of subunits B (blue), B-1 (cyan), and B-2 (gray) depicted as ribbon diagrams. Initially bound to B-1, subunit B unflattens, straining contact between subunits until they separate. The resulting loose lateral connection causes subunit B to fluctuate dynamically with respect to the rest of the filament while secured by its D-loop to subunit B-2. The simulation is aligned to $C_\alpha$ atoms of subunits B-1 and B-2, and smoothed over 3 ns. Data comes from simulation 1, and the duration is 742 ns (Table S1).

**Movie S2. The ADP-bound pointed end subunit P-1 forms unique contact with subunit P.** A movie of subunits P (red) and P-1 (pink) depicted as ribbon diagrams. The alpha-helix composed of residues 55-64 (gray) in subdomain 2 of P-1 shifts towards subunit P. Residue contacts in the hydrophobic plug (violet) of subunit P stabilize P-1 in the new large dihedral angle conformation. The D-loop of subunit P (green) also associates with subunit P, which makes the D-loop less available for binding to incoming actin monomers. The formation of the new contacts stabilizes subunit P onto the filament. Data comes from simulation 7, and the duration is 808 ns (Table S1)

**Supplemental Figures**

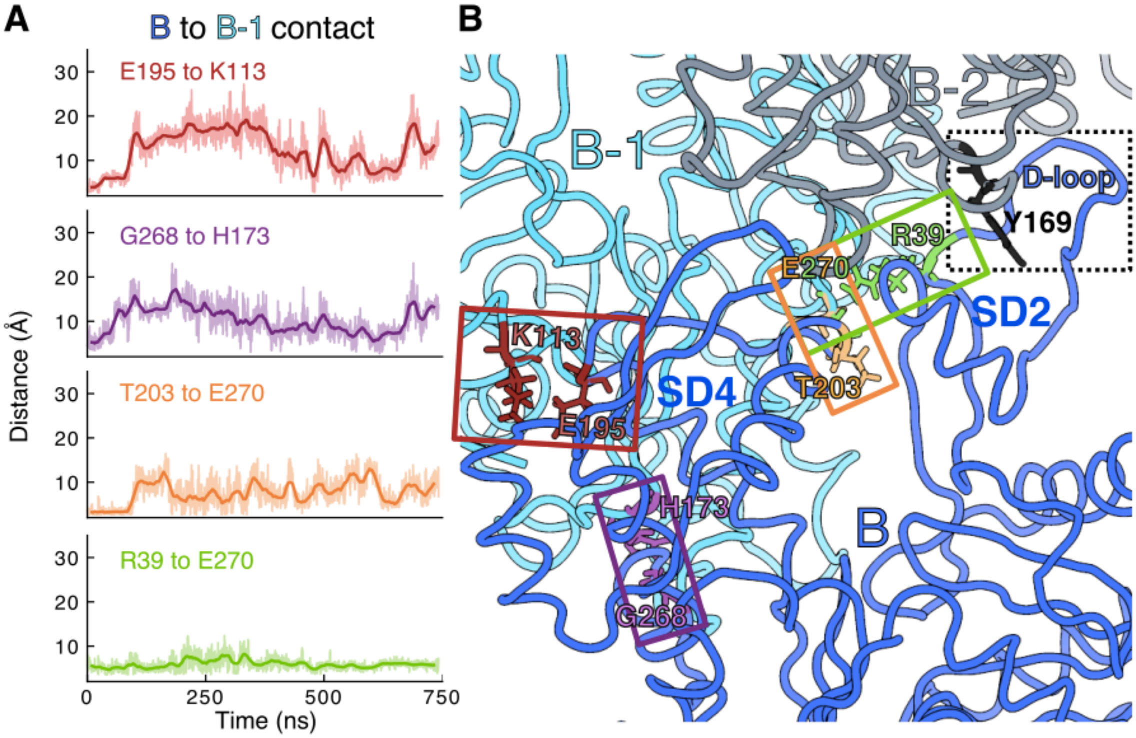


**Fig. S1. Lateral contacts between subdomain 4 of interior subunit i and subunit i-1 are broken between barbed end subunits B and B-1 in simulations of ATP and ADP-P$_i$ filaments.** (**A**) The distance between the backbone O of E195 of subunit B and the backbone N of K113 of subunit B-1 increased significantly (top, red), while the distance between the terminal sidechain carbons of R39 of subunit B and of E270 of subunit B-1 increased very little (bottom, green). The distance between the backbone O of G268 of subunit B and sidechain nitrogen NE2 of H173 of subunit B-1 (purple), as well as the distance between the backbone N of T203 of subunit B and the sidechain O (OE2) of E270 of subunit B-1 (orange) both also increased to the point of no longer mediating a stabilizing inter-subunit connection. Contact partially reform around 600 ns before being lost again. Data comes from simulation 1 (Table S1). (**B**) The contacts that break are located in subdomain 4 of subunit B (labeled SD4), whereas those that connect subdomain 2 of subunit B to subunit B-1 are preserved. This is because the D-loop of subunit B maintains a highly stable attachment to Y169 of subunit B-2 (dashed box), which steadies all of subdomain 2. Subunits B (blue), B-1 (cyan), and B-2 (gray) are depicted in ribbon representation, whereas specific contacts are depicted in licorice representation and contact pairings are colored according to the corresponding time course shown in panel A.

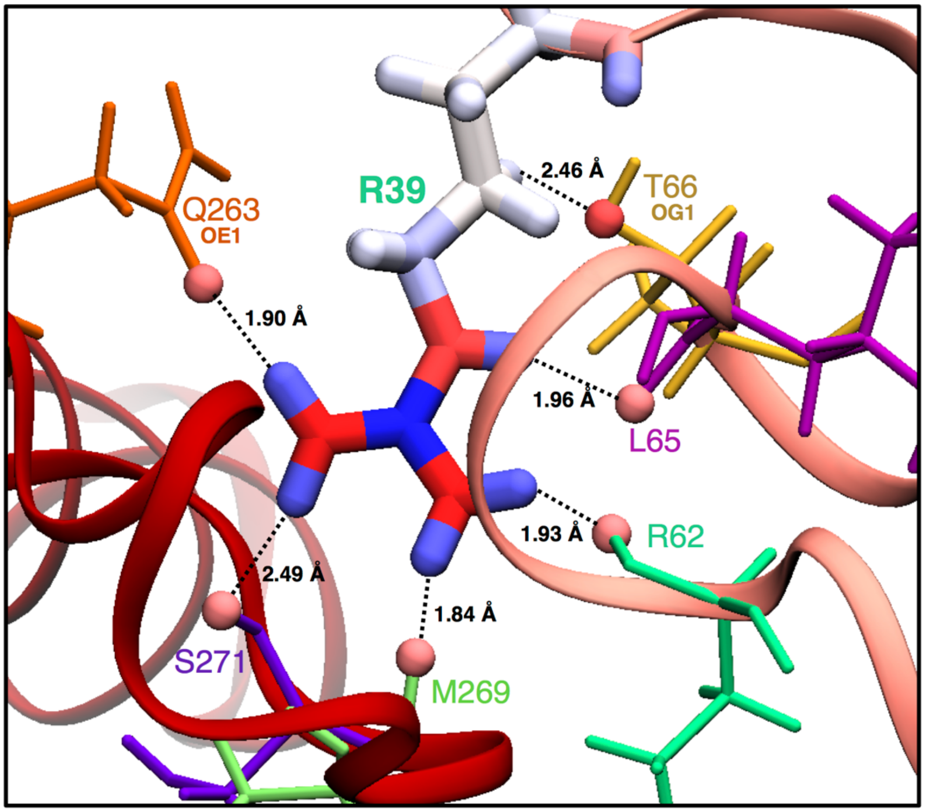


**Fig. S2. A hydrogen-bond network stabilizes R39 of subunit P-1 with oxygens of nearby residues in the connection between subunit P-1 and subunit P.** R39 of subunit P-1 is depicted in licorice representation with atoms colored by charge (blue is positive; red is negative). Subunits P-1 and P are shown as pink and red ribbons, respectively. Atoms within 2.5 Å of R39 of P-1 are shown as spheres and colored by charge; dotted lines depict the distance to atoms of R39. All atoms within the cutoff are backbone oxygens unless labeled, in which case they are sidechain oxygens. The entire residue containing the nearby oxygen atoms is depicted in a thin licorice representation colored by amino acid species. This snapshot is from the last frame of simulation 7 (Table S1)—in other frames, the backbone oxygens of G63 of subunit P-1 and E270 of subunit P participated in the hydrogen-bond network.

| Table S1: Simulations performed in this study | | | | | | |
|---|---|---|---|---|---|---|
| Number | F or G | Bound nucleotide | Replica # | Time (ns) | PDB ID | Initial D-loop |
| 1 | F | ATP | 1 | 742 | 6DJM | Extended |
| 2 | F | ATP | 2 | 338 | 6DJM | Extended |
| 3 | F | ATP | 3 | 415 | 6DJM | Extended |
| 4 | F | ADP-$P_i$ | 1 | 323 | 6DJN | Extended |
| 5 | F | ADP-$P_i$ | 2 | 798 | 6DJN | Extended |
| 6 | F | ADP-$P_i$ | 3 | 423 | 6DJN | Extended |
| 7 | F | ADP | 1 | 808 | 6DJO | Extended |
| 8 | F | ADP | 1 | 327 | N/A | Folded |
| 9 | G | ATP | 1 | 347 | 1NWK | Extended |
